# Random Sequences Rapidly Evolve into *de novo* Promoters

**DOI:** 10.1101/111880

**Authors:** Avihu H. Yona, Eric J. Alm, Jeff Gore

## Abstract

How do new promoters evolve? To follow evolution of *de novo* promoters, we put various random sequences upstream to the lac operon in *Escherichia coli* and evolved the cells in the presence of lactose. We found that a typical random sequence of ~100 bases requires only one mutation in order to enable growth on lactose by increasing resemblance to the canonical promoter motifs. We further found that ~10% of random sequences could serve as active promoters even without any period of evolutionary adaptation. Such a short mutational distance from a random sequence to an active promoter may improve evolvability yet may also lead to undesirable accidental expression. We found that across the *E. coli* genome accidental expression is minimized by avoiding codon combinations that resemble promoter motifs. Our results suggest that the promoter recognition machinery has been tuned to allow high accessibility to new promoters, and similar findings might also be observed in higher organisms or in other motif recognition machineries, like transcription factor binding sites or protein-protein interactions.

## Introduction

How new functions can evolve is a fundamental question in biology. For many complex traits a combination of genetic changes is required before a beneficial function can be obtained^1^. In such cases, the evolutionary “path” is not trivial, as the negligible selective advantage of the first mutations may prevent them from spreading in the population and further acquire the other needed mutations. The possibility of acquiring multiple desired mutations simultaneously (rather then serially) has very low probability, especially in asexual populations like bacteria that are unable to combine mutations that were acquired in different individuals^2^.

We chose bacterial promoters as a test case for the evolution of new genetic functions. The RNA polymerase requires particular sequence elements in order to transcribe a gene, and additional features like transcription factors and small ligands can further affect its activity. The *E. coli* canonical σ^70^ promoter (which is the primary sigma factor in *E. coli*) is recognized by consensus sequence elements, the two principal ones being the −10 element TATAAT and the −35 element TTGACA, which are separated by a spacer of approximately 17 bases. Additional sequence elements like the extended −10 and the UP elements can be recognized as well, and they act together for the promoter to be recognized by the RNA polymerase^3^.

The extensive study of promoters by genomic analysis^4–6^, experimental protein-DNA interactions^7–9^ and promoter libraries^10–13^ has mostly revolved around highly refined promoters i.e. long-standing wild-type promoters and their derivatives. However, the emergence of new promoters, for example when cells need to activate horizontally transferred genes^14,15^, is less understood. Recent studies have demonstrated how new promoters can emerge from duplication of existing promoters via genomic rearrangements^16,17^, transposable elements^18,19^, or by inter-species mobile elements^20^. Yet, little is known about promoters evolving *de novo*. The sequence space that encompasses the different promoter elements is extreme in size and it is unclear how far a functional promoter is from a typical random sequence. Especially in experimental and quantitative terms, the question is how many mutations does one need in order to make a functional promoter starting from a random sequence of a specific length? The mutational distance from a random sequence (composed of A, C, G and T in equal probabilities) to a functional promoter might require multiple mutations in order to have any significant level of expression. As mentioned above, in cases where multiple mutations are necessary for functionality, the evolutionary search is difficult because the first mutation does not have a selective advantage until additional mutations appear. A new promoter in such cases would probably be obtained by copying an existing promoter from elsewhere in the genome.

Exploring the fitness landscape of promoters in order to understand how non-functional sequences turn into functional promoters can be done artificially, by using pooled promoter libraries that allow the measurement of a large number of starting sequences. However, pool competition is less applicable for following an evolutionary process that requires mutational steps, as selection in pool is often dominated by a small fraction of the sequences that exhibit high activity. Therefore, to explore the fitness landscape of emerging promoters in a way that is similar to evolution in natural ecologies, we utilized lab-evolution methods. We evolved parallel populations, each starting with a different random sequence, for their ability to evolve new promoters. Following these evolving populations highlighted that new promoters can emerge from random sequences by stepwise mutations, and significantly less frequently by copying an existing promoter. Substantial promoter activity can typically be achieved by a single mutation in a 100-bases sequence, and could be further increased in a stepwise manner by additional mutations that improve similarity to the canonical promoter consensus sequences. We therefore find a remarkable flexibility of the transcription network on the one hand, with a tradeoff of low specificity on the other hand, that together raise some interesting implications on the design principles of genome evolution.

## Main Text

To create an ecological scenario that tests how bacteria can evolve *de novo* promoters, we sought a beneficial gene in the genome but not yet expressed, similarly to what might occur when a gene is transferred horizontally without a functional promoter. To this end, we chose to study evolution of a modified lac operon in *E. coli* with the native promoter replaced by random sequences. It is important to note that this work is not focused on the lac promoter or operon, as we merely use the lac metabolic genes for their ability to confer a fitness advantage upon expression in the presence of lactose. Accordingly, we modified the lac operon so that only the lac metabolic genes (*LacZYA*) remain intact (including their 5’UTR); we deleted the lac repressor (*LacI*) and eliminated the lac promoter by deleting the entire intergenic region upstream to the lac genes and replaced it with a variety of non-functional sequences. To broadly represent the non-functional sequence-space we used random sequences (generated by a computer) with equal probabilities for all four bases (Methods).

The random sequences that replaced the WT lac prompter were 103 bases long, which is a typical length for an intergenic region in *E. coli* (the median intergenic region in *E.coli* is 134 bases^21^). Also, it is the exact same length as the deleted intergenic region that originally harbored the WT lac promoter. In addition, the lactose permease (*LacY*) was fluorescently labeled with YFP^22^ for future quantification of expression. To avoid possible artifacts associated with plasmids, all modifications were made on the *E. coli* chromosome^23^, so the engineered strains had a single copy of the metabolic genes needed for lactose utilization, yet without a functional promoter (Figure 1A). We began building such strains with random sequences as intergenic regions upstream to the lac genes, and already observed for the first strains obtained that they could not utilize or grow on lactose because they could not express the lac genes. This experimental observation was therefore consistent with the expectation that a random sequence is unlikely to be a functional promoter.

**Figure 1:**
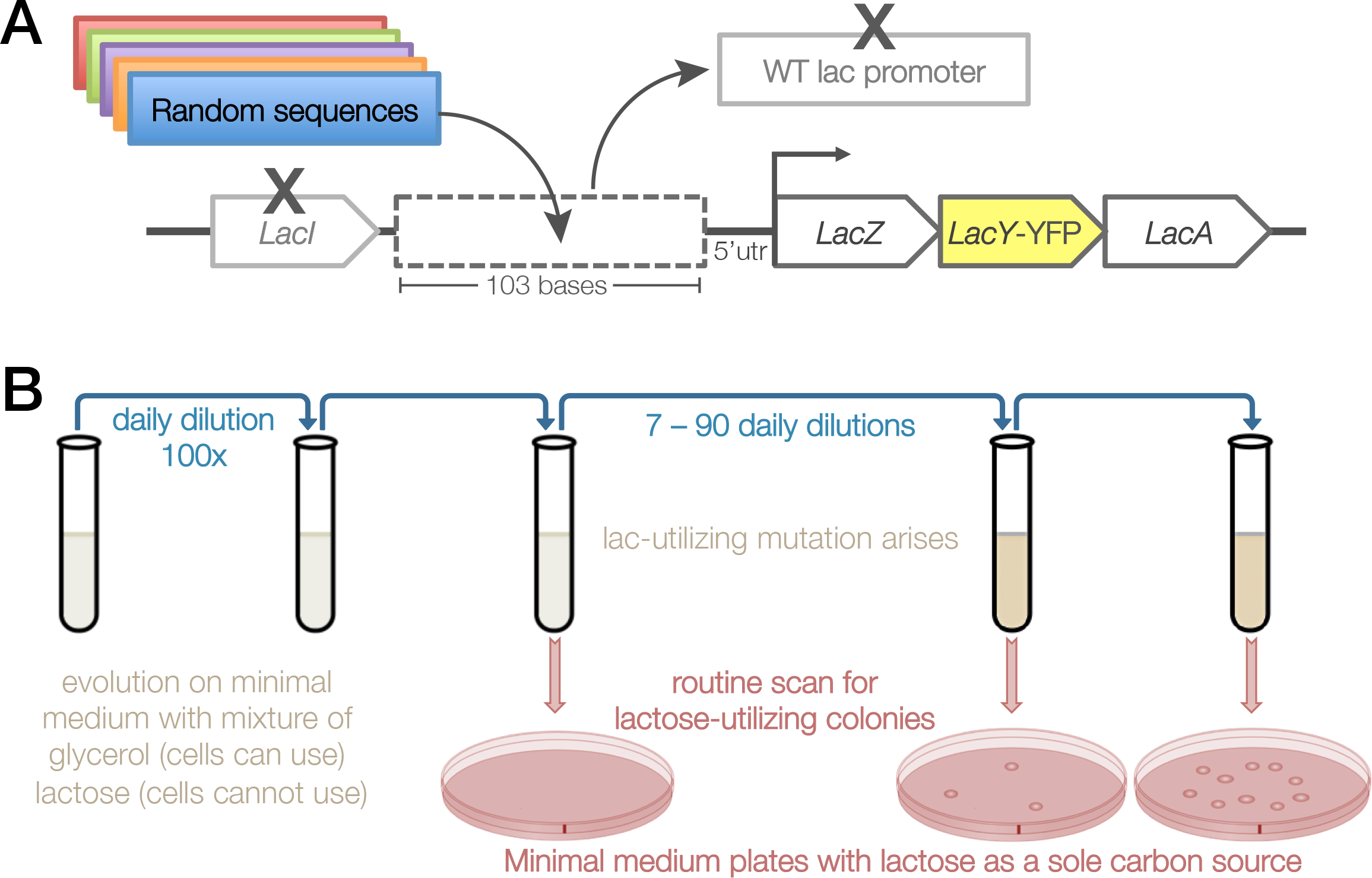
Experimental setup for evolving promoters from random sequences. **(A)** We modified the chromosomal copy of the lac operon by replacing the intergenic region that harbors the WT lac promoter with a random sequence of the same length (103 bases) that abolished the cells’ ability to utilize lactose. In addition, the lacY was tagged with YFP and the lac repressor (lacI) was deleted. **(B)** Cells were evolved by serial dilution in minimal medium containing both glycerol (0.05%), which the cells can utilize, and lactose (0.2%), which the cells cannot utilize unless they evolve de novo expression of the lac genes. During evolution, samples were routinely plated on minimal medium plates with lactose as a sole carbon source, for isolation of lactoseutilizing mutants.

To evolve *de novo* expression of the lac genes we selected for the ability to utilize lactose. Therefore, our criterion of whether expression is on or off was not by setting an arbitrary threshold, but rather by a functional readout – the ability to grow on lactose as a sole carbon source. We started evolution with a variety of strains, each one carrying a different random sequence upstream to the lac genes. We first focused on three such strains (termed RandSequence1, 2 and 3) and tested their ability to evolve expression of the lac genes, each in four replicates. As controls, we also evolved a strain in which the WT intergenic region upstream to the lac genes remained intact (termed WTpromoter), and another strain in which the entire lac operon was deleted (termed ΔLacOperon). Before the evolution experiment, only the WTpromoter strain could utilize lactose (Supp. Figure 1). Therefore, to facilitate growth to low population sizes the evolution medium contained glycerol (0.05%) that the cells can utilize and lactose (0.2%) that the cells can only exploit if they express the lac genes.

To isolate lactose-utilizing mutants, we routinely plated samples from the evolving populations on plates with lactose as the sole carbon source (M9+Lac) (Figure 1B). Remarkably, within 1-2 weeks of evolution (less than 100 generations), all of these populations exhibited lactose-utilizing abilities, except for the ΔLacOperon population (Supplementary Information). These lab evolution results therefore argue that the populations carrying random sequences instead of a promoter can rapidly evolve expression. Next, we addressed the question of whether the solutions found during evolution were mutations in the random sequences or simply copying of existing promoters from elsewhere in the genome.

To determine the molecular nature of the evolutionary adaptation, we sequenced the region upstream to the lac genes (from the beginning of the lac genes through the random sequence that replaced the WT lac promoter and up to the neighboring gene upstream). Within each of the different random sequences a single mutation was found to confer the ability to utilize lactose by *de novo* expression of the lac genes. Continued evolution yielded additional mutations within the random sequences that further increased expression from the emerging promoters. All replicates showed the same mutations, yet sometimes in different order (Supp. Table 1). Each mutation was inserted back into its relevant ancestral strain, thus confirming that the evolved ability to utilize lactose is due to the observed mutations.

Next, we assessed differences in expression by YFP measurements (thanks to the *LacY*-YFP labeling), where we found that the evolved promoters led to expression that was comparable to the fully-induced WT lac promoter (Figure 2A). This experimental evolution demonstrates how non-functional sequences can rapidly become active promoters, in a stepwise manner, by acquiring successive mutations that gradually increase expression. Next, we aimed to determine the mechanism by which these mutations induced *de novo* expression from a random sequence.

**Figure 2:**
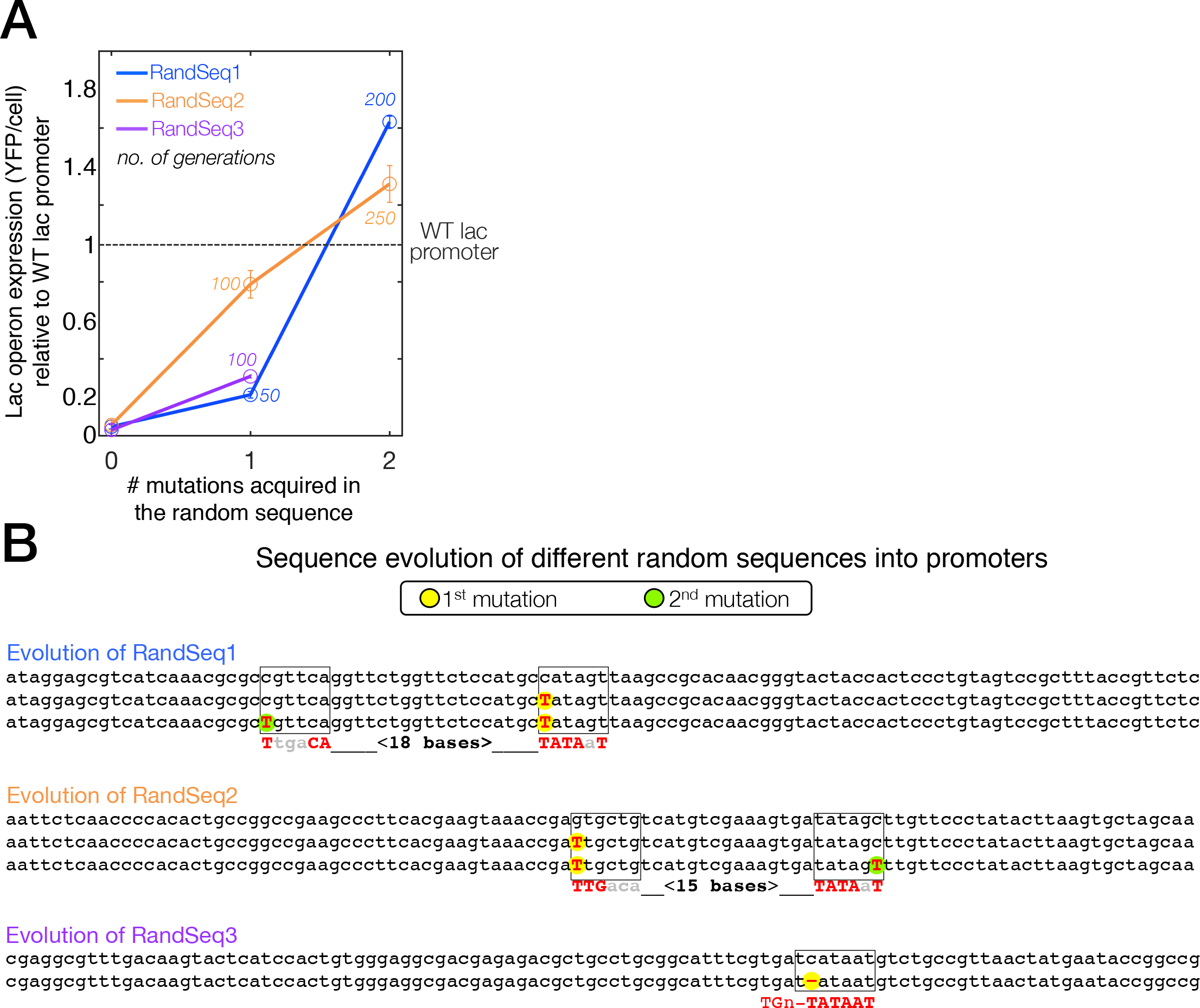
From a random sequence to an active promoter by stepwise mutations that build a canonical promoter (example from three random sequences) **(A)** Evolved expression levels of the lac genes are plotted for three strains that carry different random sequence (blue, orange and purple) as a function of the number of acquired mutations. Expression level of 1 is defined as the expression measured from the WT lac promoter, and 0 is defined as the background read of the control strain ΔLacOperon (in which the lac operon is deleted and no YFP gene was integrated). After accumulation of mutations, de novo expression is observed (as well as the ability to utilize lactose). The number of generations is indicated near each mutation. Mutations shown were verified by reinsertion into their non-evolved ancestors. **(B)** Sequences of the evolving promoters. For each strain, the top sequence is the random sequence before evolution, 2nd and 3rd lines are the random sequence with the evolved mutations (1st and 2nd mutations respectively). Increasing similarity to the canonical E. coli promoter motifs can be observed by the different mutations. For each evolving promoter the canonical promoter is shown as the bottom line where capital bases indicate a match. For RandSeq3 the evolved promoter is an extended −10 promoter, thus the −35 motif is not indicated.

The sequence context of the emerging mutations suggests that *de novo* expression has evolved by increasing similarities, in the random sequences, to the consensus sequence of the canonical promoter motifs^24^. Each of the five evolved mutations that were found in Randseq1, 2 and 3 increased the similarity to either the TATAAT or the TTGACA consensus sequences. In Randseq1 a single base substitution created an almost perfect −10 motif and a consecutive mutation further increased expression by improving the −35 element. A similar scenario was observed in Randseq2, yet in the reversed order as the first mutation created a −35 element and the later mutation further increased expression by improving the −10 element (Figure 2B). In Randseq3, however, no successive mutations were found after the first mutation that induced expression by creating a perfect TATAAT motif. The evolved mutation in Randseq3 occurred alongside an extended −10 motif^25^ that enabled expression even without a proper −35 element. Yet unlike Randseq1 and 2, in Randseq3 no putative −35 element could be found in a tolerable spacing from the −10 element. This lab-evolution experiment suggest that *de novo* promoters are highly accessible evolutionarily, as in all populations a single mutation created a promoter motif that enabled growth on lactose, suggesting that a sequence space of ~100 bases might be sufficient for evolution in order to find an active promoter with one mutational step.

The important step of random sequences evolving into functional promoters was the first mutation that was sufficient to enable growth on lactose by turning on expression. Therefore, we predicted that if indeed a single mutation in a 103-base random sequence is often sufficient to generate an active promoter, there might also be a small portion of random sequences that are already active without the need of any mutation. Indeed, when we expanded our collection to 40 strains, each carrying a different random sequence (RandSeq1 to 40), we observed that four of the strains (10%) formed colonies on M9+Lac plates before evolution and without acquiring any mutation in their random sequences. We scanned the random sequences of these already-active strains (RandSeq7, 12, 30, 34) and found regions with high similarity to the canonical promoter consensus sequences, equivalent to the similarities caused by the mutations mentioned earlier for RandSeq1, 2 and 3 (Supp. Figure 2). Given that a single mutation might be sufficient to turn expression on, we proceeded with the strains that could not grow on lactose, by putting them under selection for lactose utilization both by the abovementioned daily-dilution routine (in M9+GlyLac) and by directly screening for mutants that can form colonies on M9+Lac plates (Methods).

Overall, evolving expression of the lac operon by selection for lactose utilization was successful for all but two of the random-sequence strains (38/40). Analysis of all forty strains and their lac operon activating mutations showed that: 10±5% were already active without any mutation (4/40), 57.5±8% found mutations within the 103 bases of the random sequence (23/40), 12.5±5% found mutations in the intergenic region just upstream to the random sequence (5/40) and 15±6% utilized genomic rearrangements that relocated an existing promoter of genes found upstream to the lac genes (6/40) (Figure 3A). To confirm that transcriptional read-through from the selection gene upstream did not facilitate the emergence of *de novo* promoters, six strains were made in a marker-free manner (Methods) and showed that their ability to evolve *de novo* promoters is similar to the rest of the strains. A typical random sequence of ~100 bases is therefore not an active promoter but is frequently only one point mutation away from being an active promoter.

**Figure 3:**
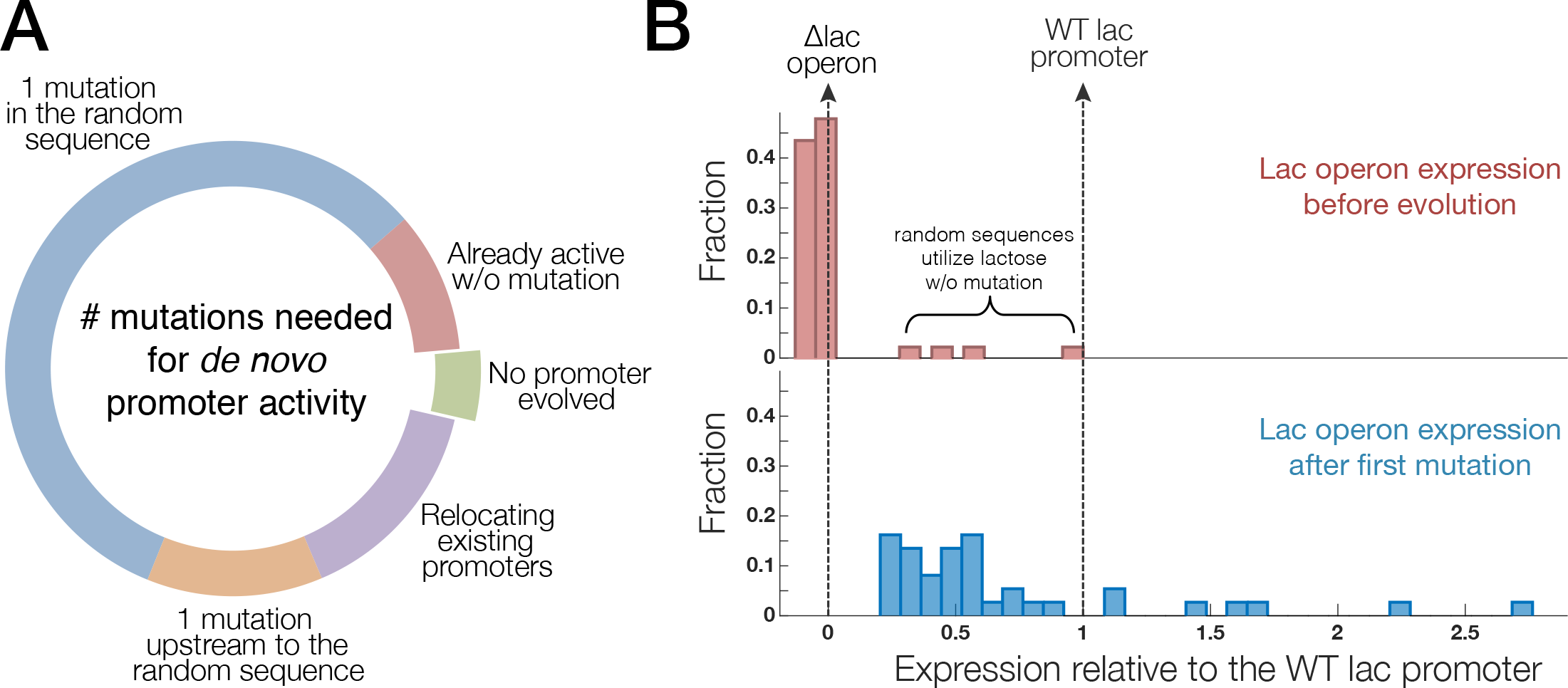
For a typical random sequence of ~100 bases one point mutation is sufficient to function as a promoter. **(A)** summary of 40 different random sequences and the different type/number of mutations by which they acquire the ability to express the lac operon and to utilize lactose. ~10% of random sequences require no mutation for such expression of the lac operon that allows growing on lactose as a sole carbon source (red segment). For 57.5% of random sequences a single mutation found within the random sequence enabled expression of the lac operon and growth on lactose (similar to RandSeq 1,2,3 shown earlier)(blue segment). Other strains either relocated an existing promoter from another locus in the genome to be upstream to the lac promoter (15%, purple) or found point mutations in the intergenic region upstream to the random sequence (12.5%, orange). **(B)** Expression of the lac operon before evolution and after the first mutation that was associated with the ability to utilize lactose (upper and lower panel respectively). Measured are YFP reads normalized to OD600 where expression level of 1 is defined as the expression measured from the WT lac promoter (right vertical dashed line), and 0 is defined as the background read of the control strain ΔLacOperon in which the lac operon is deleted and no YFP gene was integrated (left vertical dashed line). The ~10% of random sequences that conferred the ability to utilize lactose even before evolution are found to have significant expression from the lac operon (upper panel). normalized to OD600 where expression level of 1 is defined as the expression the WT lac promoter (right vertical dashed line), and 0 is defined as the background read o ain ΔLacOperon in which the lac operon is deleted and no YFP gene was integrated (left ve e). The ~10% of random sequences that conferred the ability to utilize lactose even before ound to have significant expression from the lac operon (upper panel).

YFP measurements indicated that all strains evolved substantial expression of the lac genes after acquiring the activating mutations (Figure 3B). In particular, the strains that evolved by mutations in their random sequence exhibit a median expression equivalent to ~50% of the expression observed from the fully-induced WT lac promoter (which includes a CRP transcription activator). The promoters that we evolve from random sequences therefore display significant levels of expression, and are not extremely weak “leaky” promoters. Nonetheless, continued evolution would likely lead to increased expression (as in Figure 2).

The vast majority of mutations found in the random sequence can be ascribed for increasing similarities to the two main promoter consensus sequences, the −10 and −35. Some promoters did have preexisting promoter motifs other than the −10 and −35, yet none of the mutations we found actually created or strengthened such promoter motifs, like the UP element or the TGn motif (extended −10). The “expression landscape” for promoters in this environment therefore appears to be single-peaked, as we did not observe qualitatively different sequence solutions (For details on all mutations, their verifications and different outcomes between replicates see Supp. Table 1).

Our evolution experiment showed that a single mutation could often produce expression levels similar in magnitude to the expression level obtained by the WT lac promoter. To gain a numerical perspective on these findings, we calculated the mutational distance that separates random sequences from canonical promoters of *E. coli*. To this end, we computationally created 30,000 random sequences (the same way the experimental RandSeq1 to 40 were generated) and scanned them against the canonical promoter motifs. We observed that a typical random sequence is likely to contain a promoter that captures 8 out of the 12 possible matches (of the two six-mer motifs TTGACA and TATAAT, with spacing of 17±2) (Supp. Figure 3). However, the importance of each base for promoter activity differs considerably. Essentially, three bases in each of the two motifs are highly important for promoter activity so that the core motifs of the −10 and −35 elements can often be considered as TAnnnT and TTGnnn respectively (where n means any base). We therefore re-scanned the random sequences in order to obtain the fraction of random sequences that capture, at least, these six most important bases. We found that 9±0.2% of random sequences already contain such a promoter and that 67±0.3% of them are one mutation away (Figure 4A). These “back-of-the-envelope” estimates coincide with our experimental results that showed 10±5% of random sequences that were already active and 57.5±8% that were one mutation away.

**Figure 4:**
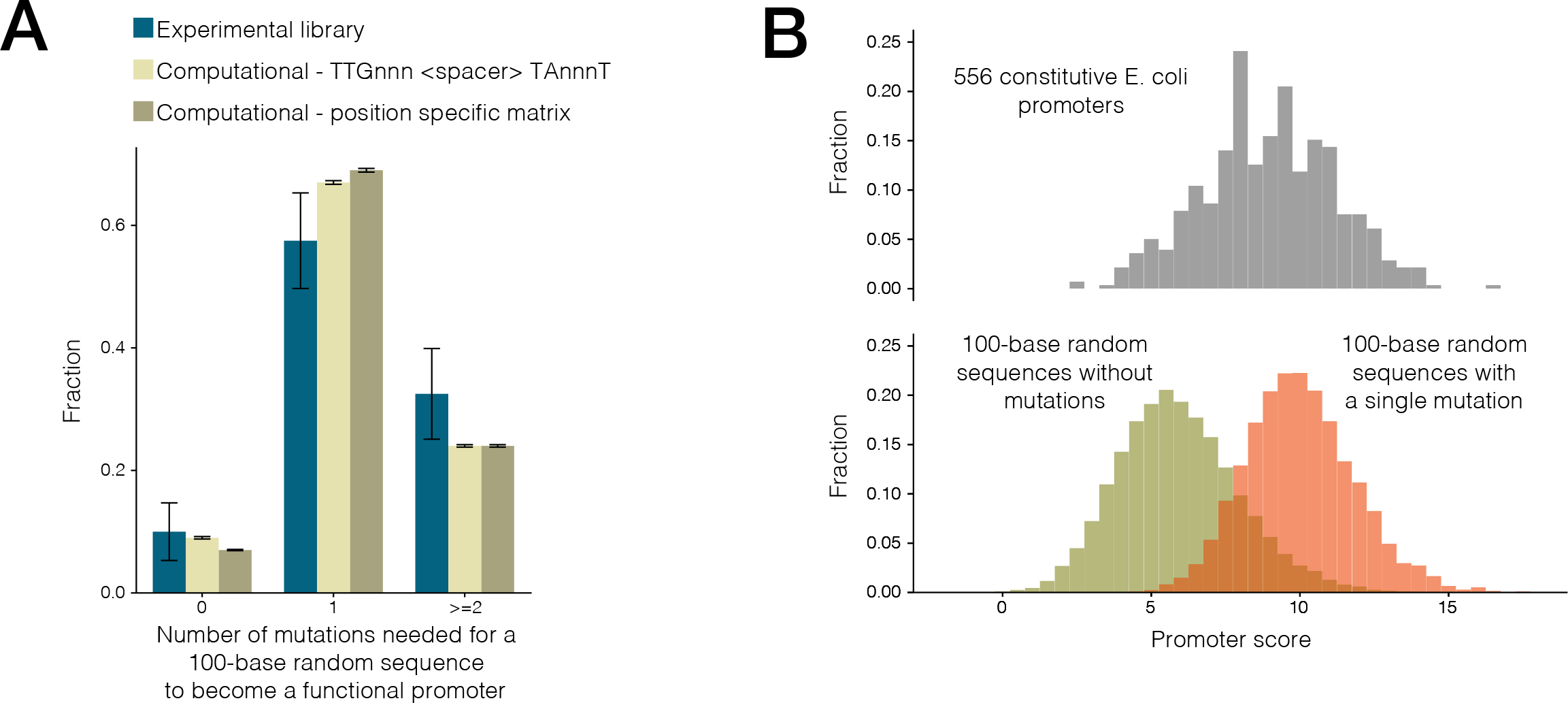
The mutational distance from a 100-base random sequence to a promoter. **(A)** The number of mutations needed to transform a random sequence to a functional promoter. The experimental results from the evolution library (n=40) are shown in blue, zero represents random sequences that were active already before any adaptation and >=2 represent random sequences that require two or more mutation in which we include all the library strains that could not evolve expression via mutations in the random sequences. Computationally generated sequences (n=30000) were scanned for their resemblance to a canonical promoter and were mutated in-silico until they pass our criterion for a promoter. We used two different criteria: the ability to capture the most important bases in the −10 and −35 promoter motifs TTGnnn and TAnnnT, with a valid spacer (light green), and the ability to score as the median score of E. coli WT constitutive promoters, according to a position specific weight matrix score (olive). Both criteria yield similar results to those of our experimental library. **(B)** Comparing promoter scores according to a position specific weight matrix. Upper panel shows a histogram of scores for E. coli constitutive promoters (n=556). Lower panel shows a histogram of scores for random generated sequences, before in-silico evolution (olive), and after the first mutation selected (orange). The overlap between the scores of the constitutive promoters to those of the random sequences (before evolution) suggests for our experimental observation of the fraction of random sequences that are already active promoters. The overlap between the scores of the constitutive promoters to those of mutated random sequences strengthen our experimental result that a random sequence of ~100 bases is typically one mutation away from functioning as a promoter. Data from these histograms were the base for the data shown in sub-figure A (olive bars).

To get a more detailed picture of the mutational distance between random sequences and active promoters, we evaluated each promoter in terms of a score that is calculated according to the hierarchical importance of the different bases that capture the canonical motifs. To this end, we weighted the bases of each promoter according to the position-specific scoring matrix of the *E. coli* canonical promoter (Methods). We performed this calculation for the random sequences mentioned above as well as for the WT constitutive promoters of *E. coli*. We found that 7±0.1% of random sequences get a higher matching score than the median score of constitutive promoters, and that 69±0.3% of random sequences need only one mutation in order to pass this score (Figure 4A). Despite the ability of a random sequence to rapidly mutate into promoters with similar matching scores to those of constitutive promoters, one should bare in mind that WT promoters also utilize additional motifs and transcription activators that may express them to higher levels than our evolved promoters. Nonetheless, our experimental result that active promoters evolve from random sequences by capturing the canonical motifs is strengthened. Indeed, a random sequence of ~100 bases typically requires only one mutation in order to reach the matching score that characterize WT constitutive promoters. Furthermore, some portion of random sequences may be active already as the matching score histograms of random sequences and constitutive promoters overlap (Figure 4B).

The short mutational distance from random sequences to active promoters may act as a double-edged sword. On the one hand, the ability to rapidly “turn on” expression may provide plasticity and high evolvability to the transcriptional network. On the other hand, this ability may also impose substantial costs, as such a promiscuous transcription machinery is prone to expression of unnecessary gene fragments ^26^. Such accidental expression is not only wasteful but can also be harmful as it may interfere with the normal expression of the genes within which it occurs ^27,28^. Our experiments indicate that ~10% of 100-base sequences can function as an active promoter, meaning that a typical ~1kb gene might naturally contain an accidental promoter inside its coding sequence. Therefore, we looked for strategies that *E. coli* might have taken to minimize accidental expression.

Normally promoters occur in the intergenic region between genes and not within the coding region, as such internal initiation of transcription can interfere with expression of the gene ^29,30^. We therefore assessed the occurrence of accidental promoters in the middle of *E. coli* genes (i.e. between the start codon of each gene till its stop codon). This coding region composes 88% of the *E. coli* genome. Since each amino acid can be encoded by multiple synonymous codons, every gene in the genome can be encoded in many alternative ways. We hypothesized that the *E. coli* genome avoids codon combinations that create promoter motifs in the middle of genes. We scanned the *E. coli* genome, looking for promoters that occur within the coding region of genes (Methods) and found that the WT *E. coli* genome has much less accidental expression than what would be expected based on a random choice of codons to encode the same amino acids, while preserving the overall codon bias (Figure 5A). The *E. coli* genome has therefore likely been under selection to avoid accidental expression within the coding region of genes.

**Figure 5:**
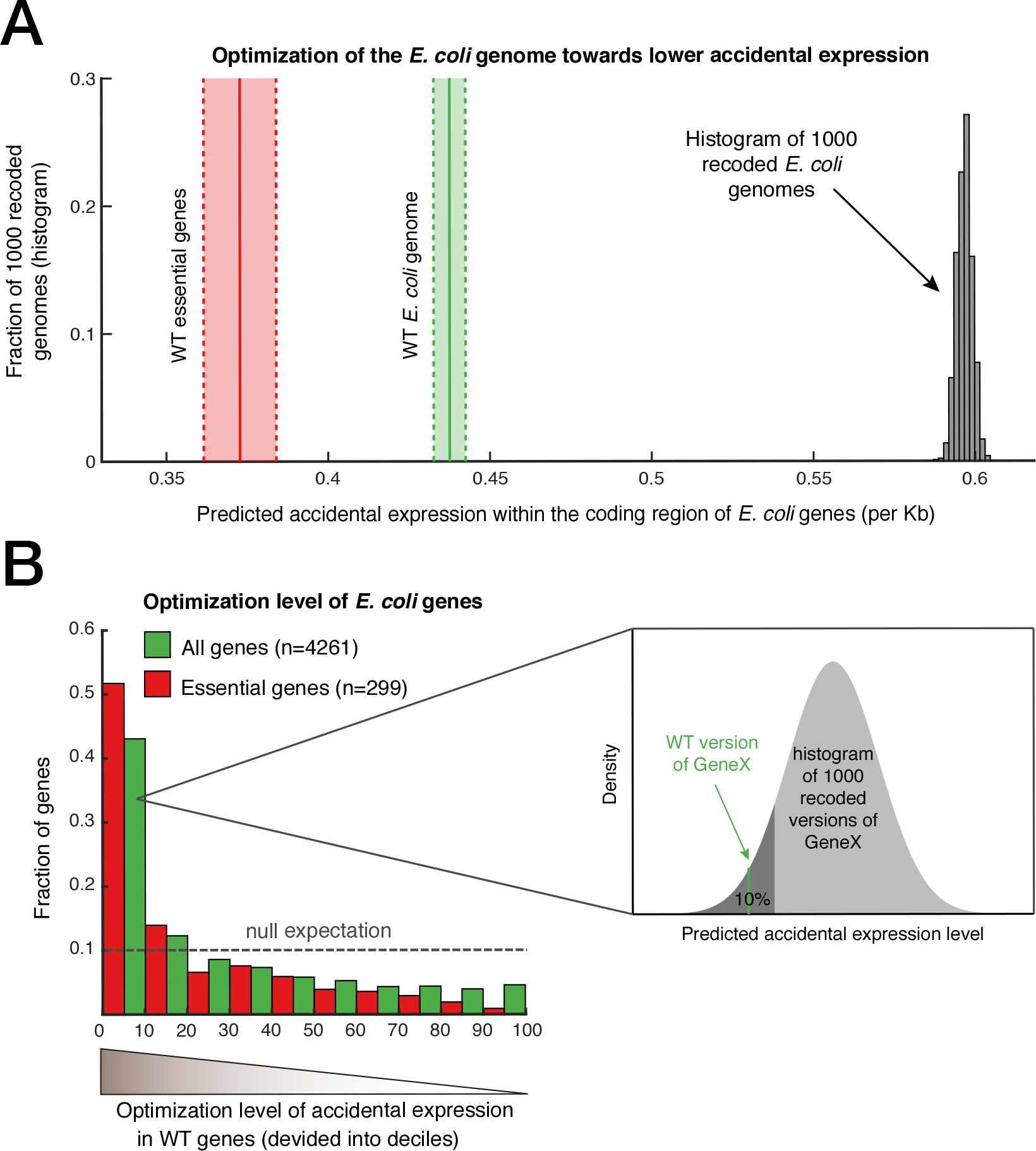
Selection against the occurrence of random promoters in the coding region of genes. We evaluated promoters that accidentally occur across the genome by searching for promoter motifs in the coding region of E. coli. As a reference we did the same evaluation for 1000 alternative versions of the E. coli coding region by recoding each gene with synonymous codons while preserving the amino acid sequence and the codon bias. **(A)** Accidental expression of the thousand recoded versions of E. coli are shown by a histogram (grey), and the accidental expression of the WT E. coli genome is shown by vertical solid lines, for all genes in green, and for the subset of essential genes in red. Shaded areas around the vertical solid lines represent S.E. (delineated by vertical dashed lines). The WT version of the genome is significantly depleted for promoter motifs, indicating genome-wide minimization of accidental expression. **(B)** For each WT gene and its 1000 recoded versions a score for accidental expression was calculated. The WT gene was then ranked in the distribution of its 1000 recoded versions (see inset illustration). Ranking values are divided into deciles, for all WT genes (green), and for the subset of essential genes (red) demonstrating that ~40% of WT genes and more than 50% of essential genes are ranked at the most optimized decile. Dashed line shows expected histogram if WT genes had similar values to their recoded version.

To assess the optimization level of each gene separately, we compared the accidental expression score of each WT gene to the scores of a thousand alternative recoded versions. Remarkably, we found that ~40% of WT genes had accidental expression as low as the lowest decile of their recoded versions. Our data indicated that some *E. coli* genes minimize accidental expression more than others. Essential genes, for example, exhibit an even stronger signal of optimization compared to the general signal obtained for all genes together (Figure 5B). Essential genes are presumably under stronger selective pressure to mitigate interference ^31,32^ and therefore they better avoid accidental expression because it leads to collisions with RNA polymerases that transcribe them ^29,30,33^. Similar results were observed when we used an alternative recoding method in which we just shuffled the codons of each gene, again indicating that the *E. coli* genome has been under selection to minimize accidental expression (Supp. Figure 4, Methods).

To further validate that the WT *E. coli* has depleted promoter motifs within its coding region, we performed a straightforward analysis by unbiased counting of six-mer occurrences across the genome. The analysis showed that promoter motifs are depleted from the middle of genes, specifically the −10 motif (Methods, Supp. Table 2). Reassuringly, among this group of depleted motifs we also found the Shine-Dalgarno 14 sequence (ribosome binding site) ^34^. Therefore, evolution may have acted to minimize accidental expression by avoiding codon combinations with similarity to promoter motifs, thereby allowing *E. coli* to benefit from flexible transcription machinery while counteracting its detrimental consequences.

## Discussion

Our study suggests that the sequence recognition of the transcription machinery is rather permissive and not restrictive^35^ to the extent that the majority of non-specific sequences are on the verge of operating as active promoters. Our experiments provide concrete quantitative measurements for this flexibility of the transcription machinery by assessing the number of mutations needed to evolve a promoter from a ~100-base random sequence, which is the characteristic length of *E. coli* intergenic regions. Specifically, we found that ~10% of random sequences need no mutation, as they are already active promoters, and that ~60% requires only a single mutation to become an active promoter. The other ~30% that evolved promoters by other means (like copying an existing promoter) or did not evolve expression at all, can be tentatively categorized as sequences that need two or more mutations for promoter functionality. We used random sequences in order to represent, in the most general way, a non-functional sequence i.e. a sequence that contains no information. It is also important to note that our assessment of whether expression is on or off does not depend on an arbitrary threshold or measurement sensitivity. The criterion for expression of the lac genes was a functional readout - the ability to grow on lactose. Furthermore, the YFP readings of the cells that evolved expression indicate that a typical new promoter obtained ~50% of the WT lac promoter expression with only one mutation (Figure 3B). This proximity of non-functional sequences to active promoters may explain part of the pervasive transcription seen in unexpected locations in bacterial genomes^26^ as well as the expression detected in large pools of plasmids that harbor degenerate sequences upstream to a reporter gene^36^.

Bacterial cells can decrease accidental expression by coiling of their chromosome, which hinders RNA polymerase from interacting with promoters, for example by histone-like proteins^37,38^. We suggest that accidental expression can also be avoided by depletion of promoter-like motifs from genomic regions that are not promoters. Specifically, codon combinations that resemble promoter motifs are depleted in genes whose expression might be sensitive to interference from internal accidental expression (like essential genes). Avoiding codon combination that resemble sequence motifs might actually be one of the constraints that have shaped the codon preferences observed in the genome. Nonetheless, accidental expression might not always be detrimental and may sometimes be selected for. When we analyzed accidental expression in toxin/antitoxin gene couples^39^, we observed higher accidental expression in toxin genes compared with their antitoxin counterparts (Supp. Figure 5, Supplementary Information). Interestingly, when we split the accidental expression score into its ‘sense’ (same strand as the gene) and ‘antisense’ (opposite strand) components, we observed that toxins had a much stronger accidental expression in their antisense direction compared to the sense direction. However, in the antitoxins, sense and antisense scores correlated, as largely seen genome-wide (Supp. Figure 6). This leads us to speculate that *E. coli* might have utilized accidental expression as a means to restrain gene expression^40,41^ of specific genes, presumably by causing head-to-head collisions of RNA polymerases^29,30,33^.

Our main findings may be relevant to other organisms and to other DNA/RNA binding proteins like transcription factors. The mutational distance between random sequences to any sequence-feature should be considered for possible “accidental recognition” and for the ability of non-functional sequences to mutate into functional ones. We demonstrated that a random sequence is likely to capture 8 out of 12 motif bases of a promoter, while natural constitutive promoters usually capture 9 out of 12. Furthermore, our experiments demonstrated that the mutational distance that separates a random sequence from a functional one, can rapidly and repeatedly found when unutilized lactose is present. Therefore, the implications of this study may also prove useful to synthetic biology designs, as one needs to be aware that spacer sequences might not always be non-functional as assumed. Moreover, spacer sequences can actually be properly designed to have lower probability for accidental functionality, for example a spacer that has particularly low chances of acting as a promoter (or ribosome binding site, or any other sequence feature).

Tuning a recognition system to be in a metastable state so that a minimal step can cause significant changes might serve as a mechanism by which cells increase their adaptability. In our study, the minimal evolutionary step (one mutation) was often sufficient to turn the transcription machinery from off to on. If two or more mutations were needed in order to create a promoter from a non-functional sequence, cells would face a much greater fitness-landscape barrier that would drastically reduce the ability to evolve *de novo* promoters. The rapid rate at which new adaptive traits appear in nature is not always anticipated and the mechanisms underlying this rapid pace are not always clear. As part of the effort to reveal such mechanisms^42^ our study suggests that the transcription machinery was tuned to be “probably approximately correct”^43^ as means to rapidly evolve *de novo* promoters. Further work will be necessary to determine whether this flexibility in transcription is also present in higher-organisms and in other recognition processes.

## Acknowledgments

We thank the Human Frontier Science Program for supporting A.H.Y. Special thanks for Idan Frumkin, Rebecca Herbst and members of the Gorelab and the Almlab for fruitful discussions. We thank the Xie lab for providing strains and Gene-Wei Li, Jean-Benoit Lalanne and Tami Lieberman for their helpful comments on the manuscript.

## Methods

**Strains** – Strains were constructed using the Lambda-Red system^23^, including integration of random sequences as promoters by using chloramphenicol resistance selection gene. Yet, for the strains with RandSeq9, 12, 15, 17, 18, 23, integration was done by the Lambda-Red-CRISPR/Cas9 system without introducing a selection marker, in order to exclude transcriptional read-through due to the expression of an upstream selection gene. The ancestral strain for all 40 random sequence strains, as well as for the control strain ΔLacOperon was SX700^22^ (also used as the control strain termed WTpromoter) in which the *lacY* was tagged with YFP. In addition, the *mutS* gene was deleted (by gentamycin resistance gene) to achieve higher yield in chromosomal integration using the lambda-red system^44^ and as a potential accelerator of evolution due to increased mutation rate. For Randseq1, 2 and 40 we created additional strains from an ancestor in which the *mutS* was not deleted and after similar evolution the exact same mutations arise. In all strains, *lacI* was deleted (for all but the CRISPR/Cas9 strains, by spectinomycin resistance gene) and replaced by an extra double terminator (BioBricks BBa_B0015) to prevent transcription read through from upstream genes.

**Random sequences** – random sequences were generated in Matlab. Each random sequence is 103 bases long, which is a typical length for an intergenic region in *E. coli* (the median intergenic region in *E.coli* is 134 bases long^21^). Also it is the same length as the WT lac intergenic region that was replaced. To prevent deviation from the overall GC content of *E. coli* (50.8%) sequences with GC context lower than 45.6% or higher than 56.0% were excluded. In addition, to avoid sequencing issues, sequences with homo-nucleotide stretches longer than five were excluded.

**Selection for lactose utilization** – Lab evolution was performed on liquid cultures grown on M9+GlyLac by daily dilution of 1:100 into 3ml of fresh medium. M9 base medium for 1L included 100uL CaCl_2_ 1M, 2ml of MgSO_4_ 1M, 10ml NH_4_Cl 2M, 200ml of M9 salts solution 5x (Sigma Aldrich). Concentrations of carbon source were 0.05% for glycerol and 0.2% for lactose for M9+GlyLac, 0.2% lactose for M9+Lac and 0.4% glycerol for M9+Gly (all in w/v). Cultures were routinely checked for increased yield at saturation and samples were plated on M9+Lac plates for isolation of colonies that can utilize lactose as a sole carbon source. In parallel to our liquid M9+GlyLac selection for lactose-utilization we also performed agar-plate selection by growing random-sequence strains on non-selective medium (M9+Gly) and then plated them while in late logarithmic phase on M9+Lac plates to select for lactose-utilizing colonies. All populations were evolved in parallel duplicates, but RandSeq1, 2, 3 had four replicates.

**Quantifying growth and expression** – Growth curves were obtained by 24h measurements of OD_600_ every 10min. Expression of the lac genes was quantified by YFP florescence measurements. Both measurements performed by a Tecan M200 plate reader. The expression of evolved cells was quantified by comparison to the control strain WTpromoter. All strains were measured for expression of the lac genes by YFP florescence by growth on M9+Gly, except WTpromoter that was grown on M9+Lac for induction of the WT lac operon.

***E. coli* genomic data** – Lists of essential genes and prophage genes were downloaded from EcoGene^21^, a list of toxin-antitoxin gene couples was obtained from Ecocyc^39^, coding sequences of genes were downloaded from GeneBank (K-12 substr. MG1655, U00096).

**Recoding the coding sequence of *E. coli* genes** – To create alternative versions of the coding region we recoded all translated genes in *E. coli* (n=4261) 1000 different times while preserving the amino acid sequence and codon bias^45^. As another null model we also shuffled the codons of each gene in 1000 permutations. Although a shuffled version of a gene does not preserve the amino acid sequence, it exactly preserves the GC content of each gene, and thus it controls for another aspect that may result in accidental expression.

**Promoter scores and prediction** – For evaluating promoters in random sequences we counted the number of matches to the canonical promoter motifs or to their core bases (TTGnnn and TAnnnT) by scanning sequences using a sliding window that identified promoter motifs with maximal agreement to the canonical *E. coli* σ^70^ promoter. When we further calculated the promoter score according to specific position weight matrix, we used a weight matrix^24^ that contains a weight for each base in the −10 and −35 elements, including a weight for the spacer length. For evaluation of accidental expression from the coding region of *E. coli* and its recoded versions we used the output from BPROM^46,47^ which takes into account all sequence motifs that affect expression (not only the −10 and −35). We obtained predicted expression scores by combining the output scores and factoring in the prediction score (LDF) from the output by multiplying.

**Six-mer analysis** – Looking for depleted and over represented motifs we counted the occurrences of all six-mers within the coding region of *E. coli*. We compiled a list of all 4096 possible six-mers and counted how many times each six-mer occurs in all WT coding region compared to the 1000 recoded versions. Then, we focused on six-mers that are significantly rare/abundant in WT version compared with their counting in the recoded versions.

## Supplementary Figure 1

**Supplementary Figure 1:**
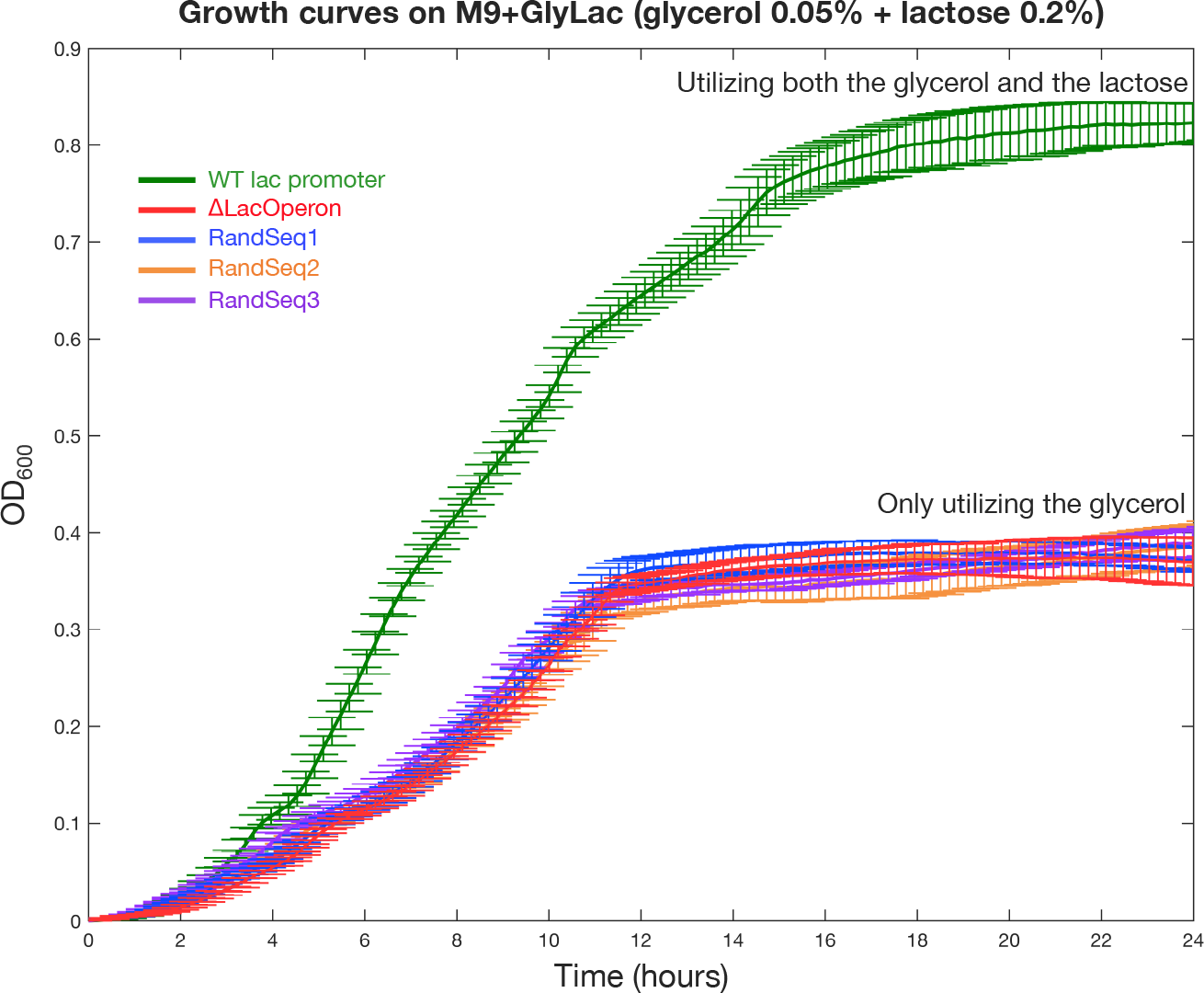
Replacing the WT lac promoter with a random sequence typically abolishes the ability to utilize lactose. Growth curve measurements of WTpromoter (green), ΔLacOperon (red) and RandSequence1, 2, 3 (blue, orange and purple respectively). Shown in values of optical density (OD600) over time during continuous growth on minimal medium (M9+GlyLac, glycerol 0.05% plus lactose 0.2%) at 37°C. The random sequence strains can only utilize the glycerol in the medium and show a growth curve very similar to the Δ LacOperon strain in which the lac genes were deleted. The difference in growth curves between the random sequence strains to WTpromoter reflects the adaptive potential for de novo expression of the lac operon.

## Supplementary Figure 2

**Supplementary Figure 2:**
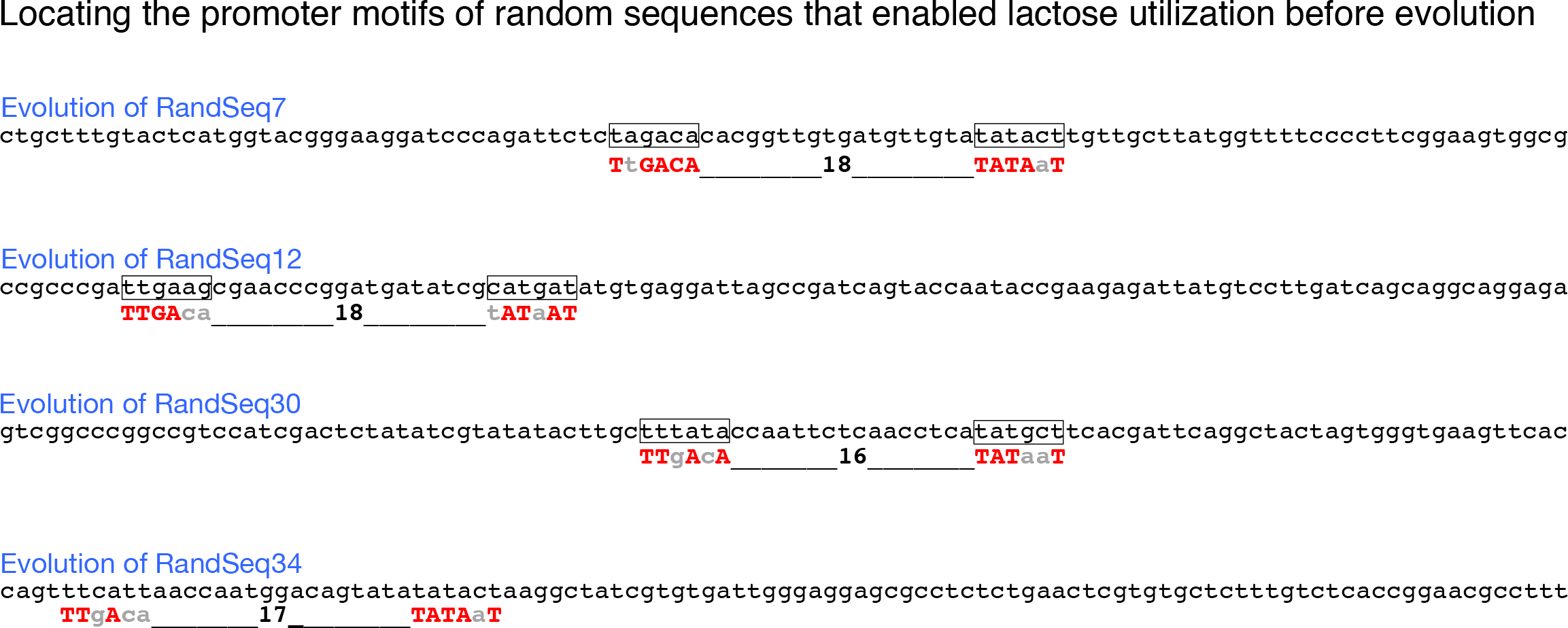
Realizing promoter motifs in the random sequences that were already active promoters before evolution. Shown are the sequences of RandSequences7, 12, 30, 34 and the locations of promoter motifs in the random sequences. For these four strains, we observed the ability of cells to grow on lactose-only plates (M9+Lac) without any adaptation. Below each random sequence the canonical promoter is shown where capital bases indicate a match to the canonical motifs TTGACA and TATAAT.

## Supplementary Figure 3

**Supplementary Figure 3:**
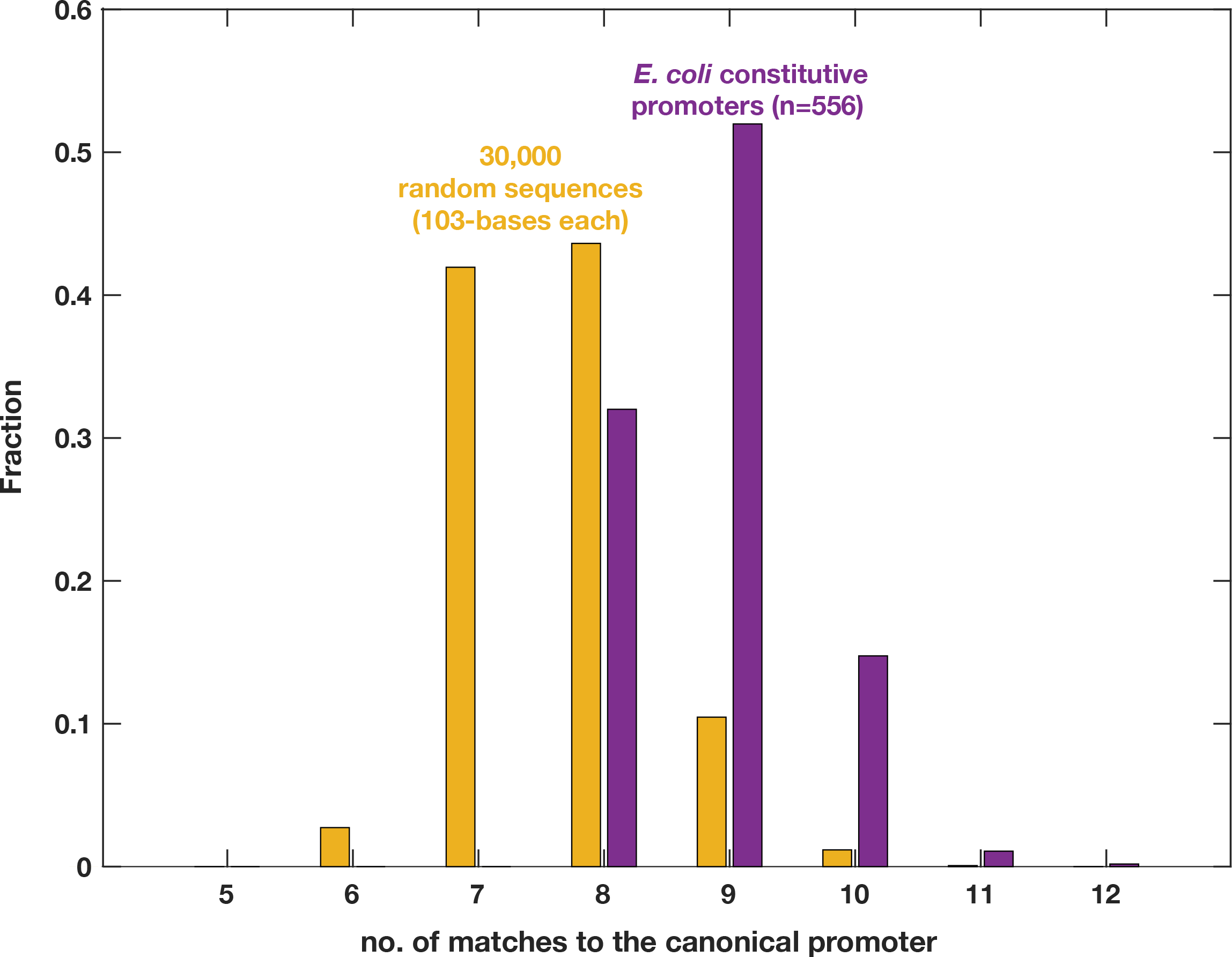
Mutational distances to the canonical promoter - random sequences are one mutational step behind E. coli constitutive promoters. The distribution of the number of matches (regardless of hierarchical importance) to the canonical promoter (defined as TTGACA, a spacer of 17±2 bases, and TATAAT) is shown for 30,000 random sequences (103 bases each) (orange), alongside the matches found for the 556 E. coli constitutive promoters (purple). The one mutation shift that separates the two distributions suggests that for many of the random sequences a single mutation can increase the number of matches to the number that characterize constitutive promoters in E. coli.

## Supplementary Figure 4

**Supplementary Figure 4:**
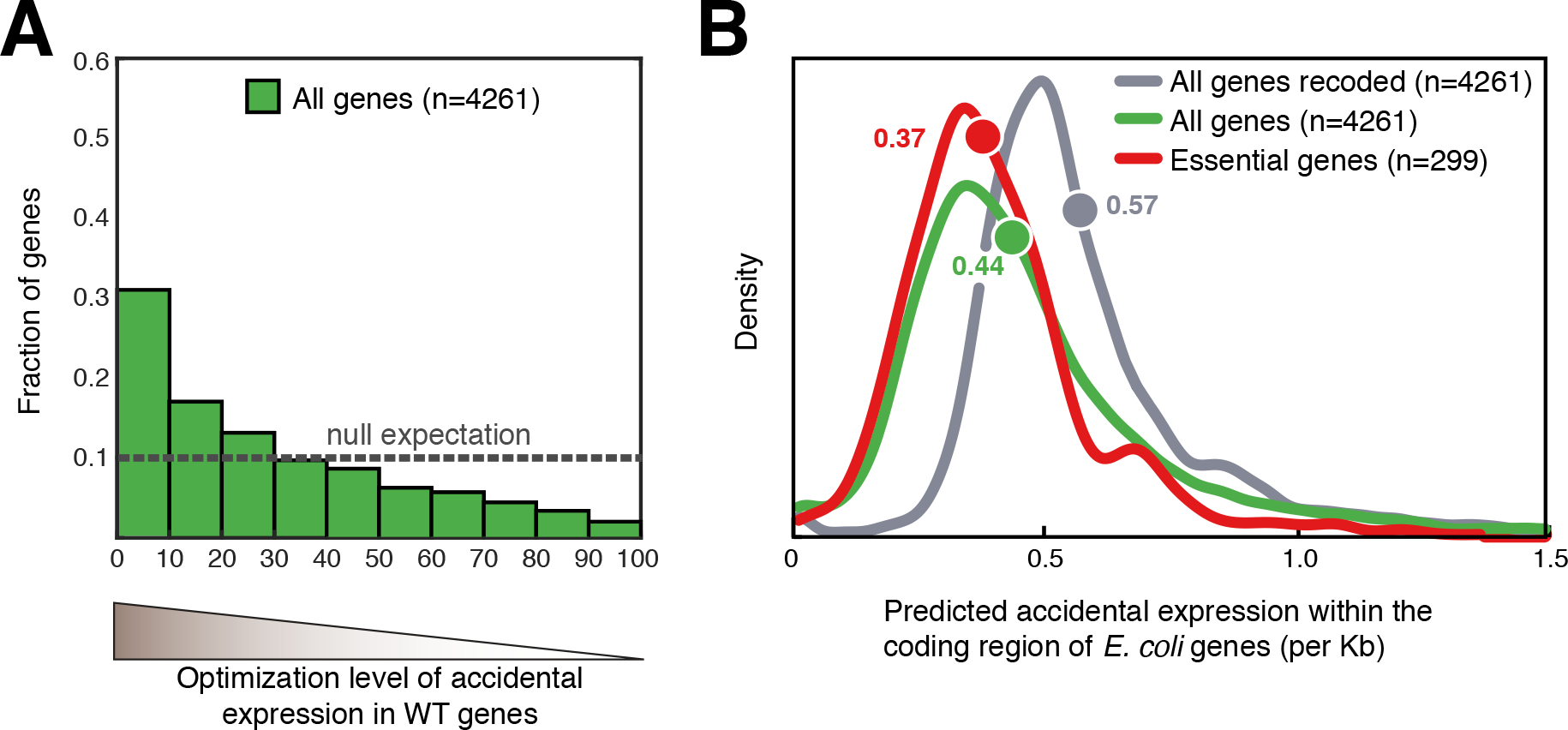
Selection against the occurrence of random promoters in the genome – alternative null model. We evaluated promoters that accidentally occur across the genome by searching for promoter motifs in the coding region of E. coli. As a reference we did the same evaluation for 1000 alternative versions of the E. coli coding region by shuffling the codons of each gene, which maintains the GC content and codon bias of each gene. Comparing the WT genes to the 1000 shuffled versions allowed us to look for codon combinations that might have been under negative selection in the WT genome. For example, the shuffled versions can indicate if a combination of two specific codons is avoided in the WT genes because it creates a promoter motif inside a gene. **(A)** A score for accidental expression is calculated for each WT gene and a rank is assigned to each gene by its order in the scores of its 1000 shuffled versions. Shown is the histogram of ranks (divided into deciles) for all WT genes demonstrating that ~30% of WT genes are ranked at the most optimized decile. Dashed line shows expected histogram if WT genes had similar values to their shuffled versions. **(B)** Density plots of accidental expression in the coding sequences of E. coli genes. Distribution of a thousand shuffled versions of E. coli coding region are shown in grey (the value that represent each gene is the median of its 1000 shuffled versions), the accidental expression of the WT E. coli genes is shown in green, and for the subset of essential genes in red. The WT version of the genome is significantly more depleted for promoter motifs, indicating genome-wide minimization of accidental expression. This minimization is further emphasized for the essential genes.genes in red. The WT version of the genome is significantly more depleted for promoter motifs, indicating genome-wide minimization of accidental expression. This minimization is further for the essential genes.

## Supplementary Figure 5

**Supplementary Figure 5:**
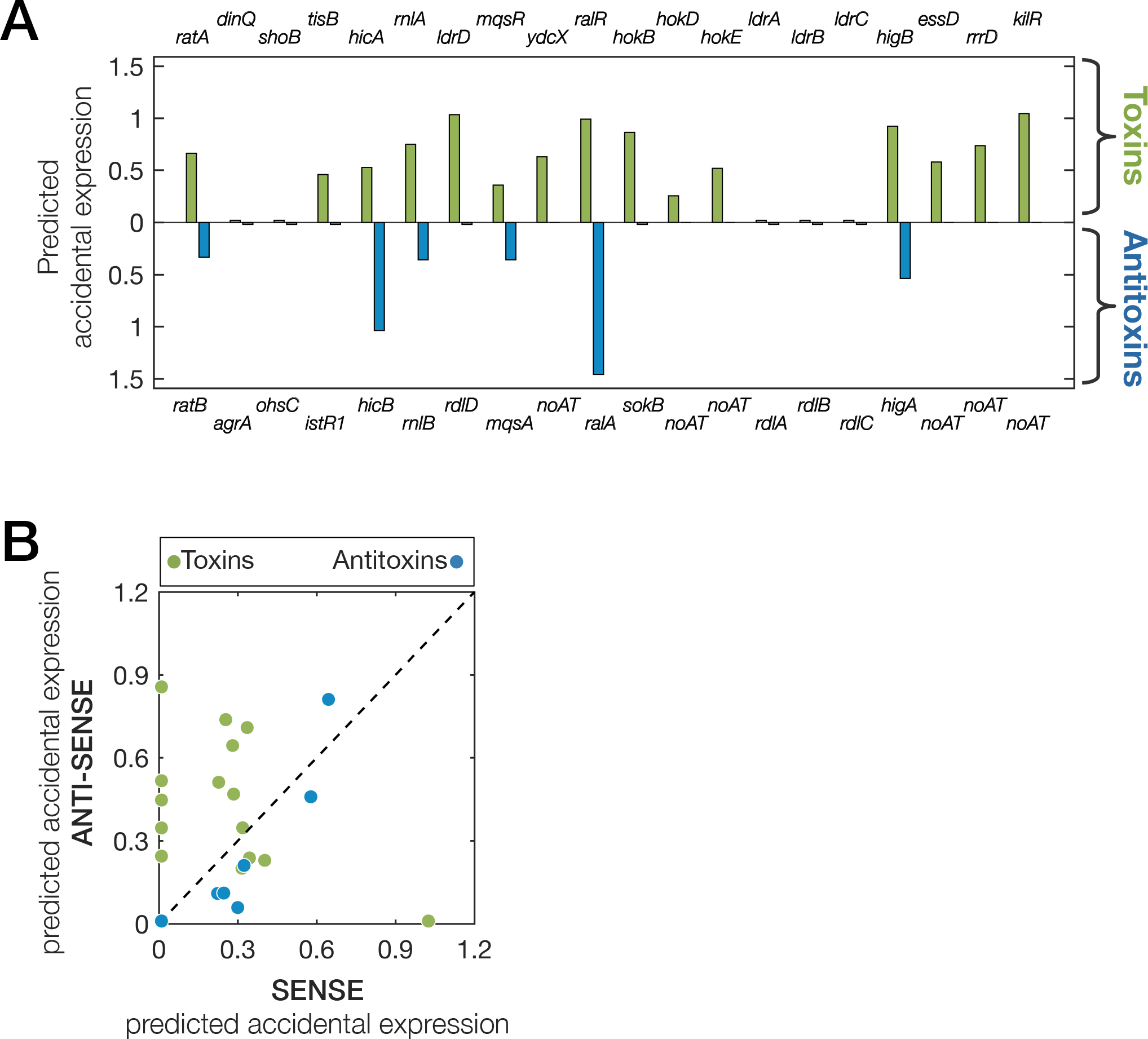
Accidental expression within toxin genes might be selected for as a means to control their expression. For each toxin-antitoxin couple the accidental expression scores examined for differences between toxins genes to their antitoxins and between accidental expressions in ‘sense’ (with the direction of the gene) compared to the ‘antisense’ (against the direction of the gene). **(A)** Accidental expression scores are compared between toxins (above the X-axis) and their antitoxin (below the X-axis) showing a tendency of toxins to have higher accidental expression compared with their antitoxin counterparts. **(B)** For both toxin and antitoxin genes the accidental expression was split into ‘sense’ and ‘antisense’ direction. While in antitoxin genes the two components tend to correlate (as generally seen in the genome, see Supplementary Figure 6) in the toxins genes the ‘antisense’ direction is significantly higher, which may imply that E. coli selects for maintaining ‘antisense’ accidental expression in order to control expression of genes whose higher dosages may harm the cells. Mechanistically, this is presumably due to the fact that antisense transcription collides with the RNA polymerase that expresses the toxin genes.

## Supplementary Figure 6

**Supplementary Figure 6:**
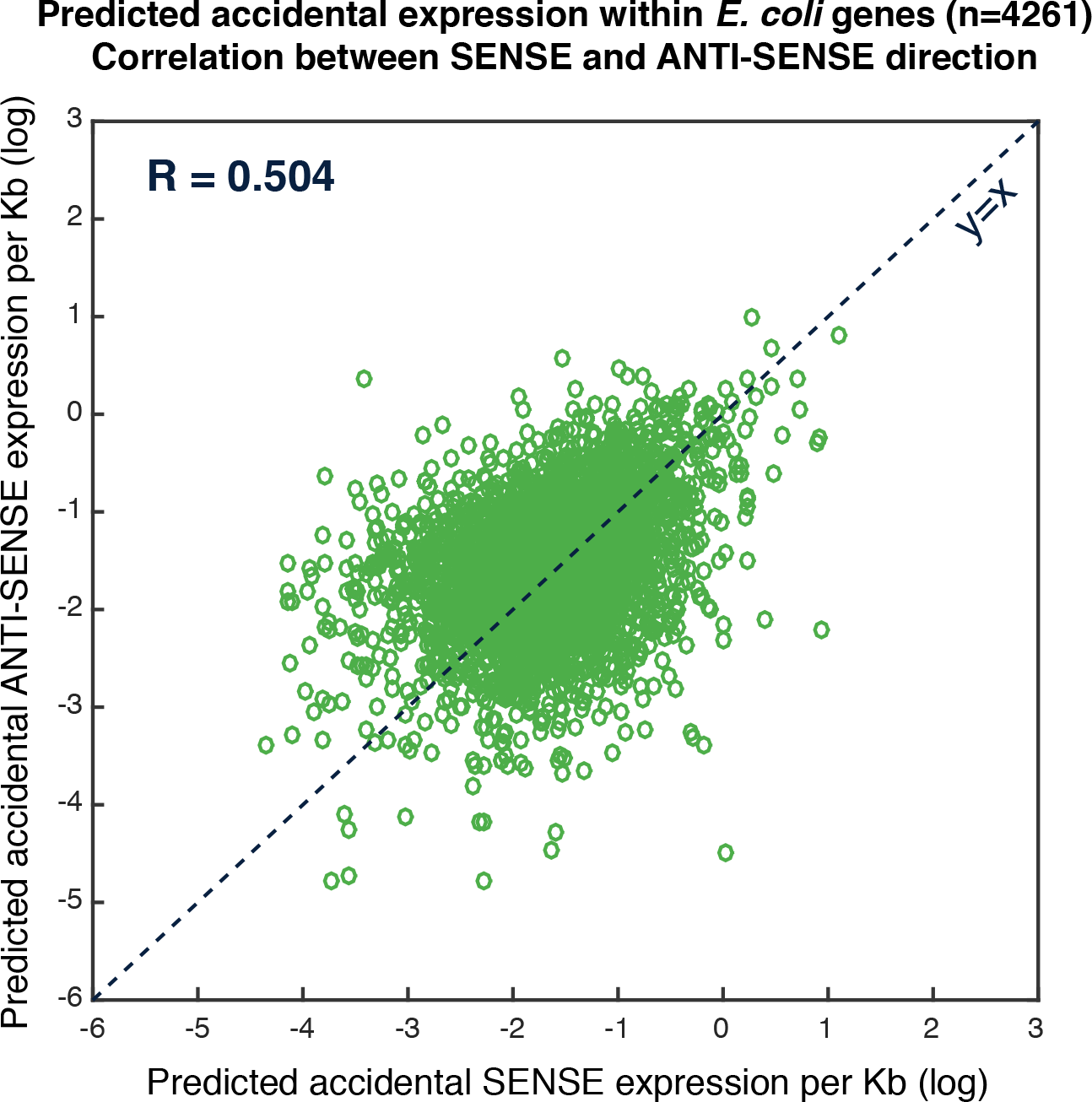
Genome-wide correlation between predicted accidental expression in ‘sense’ and ‘anti-sense’ directions. For each WT gene of E. coli we split the score obtained for accidental expression into its two contributing directions (each gene is represented by a green circle). A general correlation (R=0.504) is observed between ‘sense’ and ‘anti-sense’ directions.

## Supplementary Information

### The possibility of evolving lactose utilizing capabilities w/o the lac genes

The fact that the ΔLacOperon strain did not evolve lactose utilizing capabilities indicates that in the random sequence strains lactose utilization arose due to actual activation of the lac genes, by the verified mutations, rather than due to dubious trans–acting mutations. Furthermore, the possibility of activating an *EBG* gene^48^ (evolved β-galactosidase) is unlikely as it can only cover for the lack of *lacZ*, but still there is no active permease to replace the function of *lacY*.

### Expression activation by capturing an existing promoter or a mutation in the intergenic region upstream to the random sequence

For the random sequences listed in Extended Data Table.1 as evolved by capturing an existing promoter upstream, we observed various deletions in the intergenic region upstream to the lac genes. All of these deletions placed the lac genes in front of the upstream chloramphenicol selection gene. These deletions also eliminated the termination sequences that separated the lac genes from the genes upstream.

In strains where activating mutations appeared in the intergenic region, just upstream to the random sequence, *de novo* promoters were observed in some cases by mutations, in a similar manner to mutations that created *de novo* promoters in the random sequences (detailed in the mutations table). Yet, there was a group of point mutations, all at the same nucleotide, that occurred within the spacer of a predicted promoter that was experimentally inactive. Nevertheless, one of these mutations was sufficient for expression of the lac genes. The sequence of the predicted, yet inactive, promoter located in the intergenic region was (tcgaaa)gactgggcctttcg(ttttat), where the minus-35 site is TcGAaA and the minus-10 site is TtTtAT. This promoter has a 14-base spacer and the ‘g’ in the middle of this spacer was mutated multiple times in different strain. In some cases from g to T, in other cases from g to A and once the g was deleted (1 base deletion). It is not clear what was the mechanism by which these mutations activate expression.

We hypothesize that random sequences that evolved expression via mutations in the intergenic region might do so because they could not find an activating mutation in the random sequence. For such sequences, a mutation in the random sequence that can induce expression might not exist. Therefore, we took such a sequence, RandSeq27, and computed mutations that might improve its chances of becoming an active promoter. To this end, we scanned the original RandSeq27 for maximal matches to the canonical promoter. Since there were multiple matches, we chose the maximal match with an optimal spacer of 17 bases. Then, we introduce a point mutation that improved the minus-10 motif. After introducing this mutation into RandSeq27, it did not show promoter activity, yet after applying selection for growth on lactose (like in the library of ransom sequences) the strain found a second mutation that together with the first one we inserted exhibited expression of the lac genes:

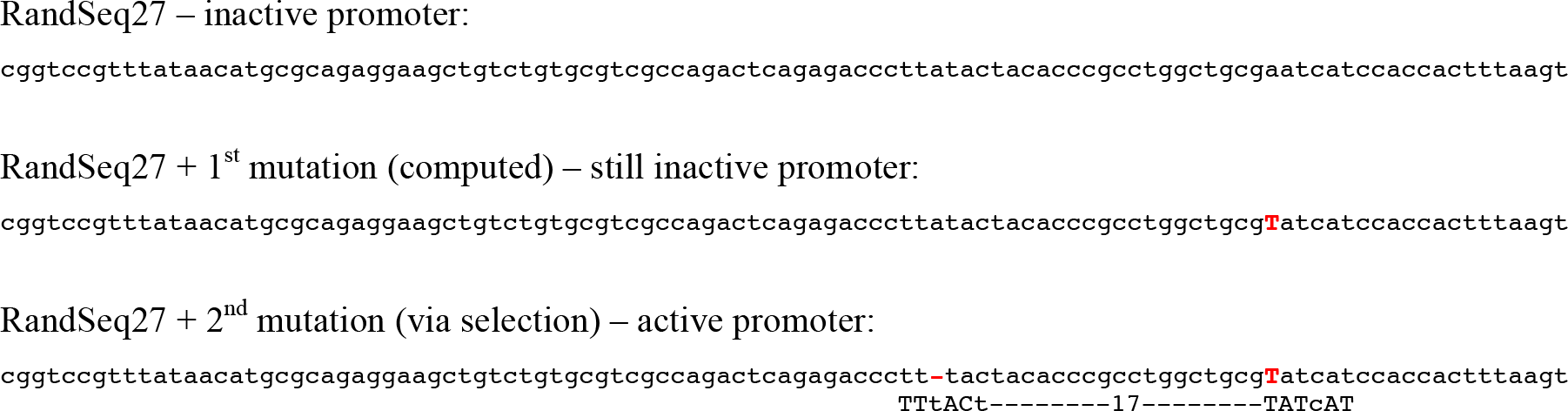

This might imply that such sequences are two mutations away from functioning as active promoters.

### The different costs of accidental expression and the motivation to focus on toxin-antitoxin gene couples

Accidental expression has a global cost due to waste of resources and occupying cellular machineries. In addition there is also a cost that is due to interference of specific genes. We observed that depletion of accidental expression is more emphasized in essential genes and is less observed in foreign genes like toxin and antitoxin prophage genes. Besides the stronger selective pressure to mitigate interference in essential genes, additional possible reasons for these differences may include: (a) foreign genes have been in the *E. coli* genome for shorter time and thus their expected optimization level is lower, and (b) foreign genes may have lower GC content than *E. coli*, which may affect accidental expression^49^ as promoter motifs are AT-rich. To decipher between these potential factors, we therefore focused on toxin/anti-toxin gene couples^39^, as for each couple the age in the *E. coli* genome is presumably the same, and they have similar GC content. Nonetheless, the anti-toxin gene is more important to the *E. coli* fitness than its toxin counterpart. Indeed, we observed lower accidental expression in anti-toxin genes compared with toxin genes. This result implies that for each gene the level of avoiding accidental expression is mainly dependent on how important to the fitness it is to have this gene expressed without interference.

### Toxin Anti-toxin couples

When analyzing toxin-antitixin gene couples for potential differences in their accidental expression, especially between sense and anti-sense orientations, we excluded gene couples whose orientation in the genome could not lead us to meaningful conclusions. Specifically, we excluded gene couples for the following reasons:

a. Toxin and antitoxin genes were overlapping, hence internal expression affects both (e.g. *ibsA* nad *sibA*).
b. Couples that had this orientation Antitoxin ➔ Toxin ➔ in which antisense expression from within the toxin gene also influences the adjacent upstream antitoxin (e.g. *yafQ* and *dinJ*).
c. Couples where the annotated promoter of the antitoxin gene is within the toxin gene and thus interference to the toxin is from a canonical functional promoter (e.g. *symE* and *symR*).

